# Evolution of *Daphnia* population dynamics following invasion by a non-native predator

**DOI:** 10.1101/2022.01.20.477096

**Authors:** Sigurd Einum, Emil R. Ullern, Matthew Walsh, Tim Burton

**Affiliations:** Centre for Biodiversity Dynamics, Department of Biology, Norwegian Univ. of Science and Technology, Høgskoleringen 5, Realfagbygget, NTNU, NO-7491 Trondheim, Norway; Department of Biology, University of Texas at Arlington, Arlington, Texas, 76019, USA; Norwegian Institute for Nature Research, Høgskoleringen 9, NO-7034 Trondheim, Norway

**Keywords:** zooplankton, density-dependent selection, predator-prey interactions, invasion ecology, bottom-up, top-down

## Abstract

1. Predators are frequently observed to cause evolutionary responses in prey phenotypes, which may, in turn, translate into evolutionary shifts in prey population dynamics. Although a link between predation and population growth has been demonstrated in experimental evolution studies, insights from natural populations are lacking.
2. Here we tested for evolutionary changes in the population dynamics of the herbivorous water flea *Daphnia pulicaria* in response to the invasion of the predatory spiny water flea (*Bythotrephes longimanus*) in the Great Lakes region, USA. Using a resurrection ecological approach and a 3-month population growth experiment (in the absence of predation) we compared population dynamics in daphnia from pre- and post-invasion time periods.
3. Post-invasion daphnia were able to maintain an overall higher population abundance throughout the growth experiment, both in terms of the number of individuals (28% higher) and total population biomass (33% higher). Estimation of population dynamics parameters from a theta-logistic model suggested that this was achieved through an increase in intrinsic population growth rate as well as increased carrying capacity.
4. The observed difference in intrinsic rate of increase could not be predicted based on previous measurements of life-history traits in these clones. This indicates that care should be taken when extrapolating from a few life history traits measured in isolated individuals under controlled conditions to population dynamics.
5. Whereas previous experimental evolution studies of predator-prey interactions have demonstrated that genotypes that have evolved under predation have inferior population growth when the predator is absent, this was not the case for the *Daphnia*. We suggest that complexities in ecological interactions of natural ecosystems, such as the potential for spatial and temporal avoidance of predation, makes it challenging to provide general predictions about evolutionary responses in population dynamics to predators.

## Introduction

Predation is an important ecological process that structures communities through numerical effects on their prey and with potentially cascading effects on lower trophic levels. However, a full understanding of predation effects likely requires knowledge about how predators influence the phenotypes of their prey. One relevant aspect in this context is that predators may influence the distribution of phenotypic traits in prey populations through selective predation and resulting evolutionary responses in targeted traits (Åbjörnsson et al., 2004; Barata et al., 2001; Fisk et al., 2007; Handelsman et al., 2013; Melotto et al., 2020). Body size and age are two of the traits that may commonly be under such selection. Thus, a change in the intensity of predation may lead to an evolutionary response in important life history traits such as age at maturation and reproductive investment (Reznick et al., 2001; Spitze, 1991). However, it has largely been overlooked that evolutionary responses in prey may also arise as an indirect byproduct of predator-induced mortality. Such selection mediated by the indirect effects of predation may stem from large-scale shifts in community structure or ecosystem function. For example, a shift in the direction of trophic control in the environment inhabited by the affected prey species may occur, from a state of ‘bottom-up control’ (i.e. resource limitation) to ‘top-down control’ (i.e. predator control), or *vice versa*. The resulting evolutionary responses in traits such as age at maturation, size and number of offspring, and competitive abilities may then be predicted based on density-dependent selection theory (Einum et al., 2008; Joshi & Mueller, 1996; Joshi et al., 2001; Mueller et al., 1991; Sæther et al., 2016). Although prey evolutionary responses resulting from indirect selection maybe harder to detect and quantify, preliminary evidence indicates that they may be of similar importance to those resulting from the direct effects of predator induced mortality on prey phenotypes (Schmitz et al., 1997; Walsh & Reznick, 2010).

Given that predators may impose direct and indirect selection on their prey and drive evolutionary changes in their phenotypes, it follows logically that this could translate into evolution of the prey population dynamics. Such changes might include shifts in the intrinsic rate of increase and/or the strength of density dependence. Evolutionary responses in prey population dynamics have been demonstrated by rearing prey under different levels of predation under laboratory conditions (Shertzer et al., 2002; Turcotte et al., 2011; Yoshida et al., 2003, 2004). However, complimentary studies of evolved responses in natural populations are scarce (Walsh et al. 2012). One reason for this is that this question can rarely be addressed by observing such phenomena in the wild, as ecological and evolutionary effects of the predator on population dynamics occur simultaneously and are confounded. Ellner et al. (2011) suggested an approach to disentangle the contribution from ecological and evolutionary effects in shaping changes in phenotypes of natural populations, but only for populations where pedigree information is available. An alternative approach is to quantify genetically based differences in life-history traits (e.g. age at maturation, age-specific fecundity) in a common environment using individuals originating from different predation regimes, and then use these to infer an intrinsic rate of population growth (i.e. *r*, Gillis & Walsh, 2017). Critically though, such data are typically obtained under *ad libitum* food availability and in the absence of density effects, meaning that a ‘true’ ecological context will be lacking in most cases.

The most dramatic changes in predation pressure may be expected when a novel predator successfully invades a new ecosystem. Invasive predators can have pervasive effects on native prey species, often causing rapid changes in both population abundance and phenotypic traits (Carroll et al., 2003; Reznick & Ghalambor, 2001; Strauss et al., 2006; Thompson, 1998). The effect of predation has received particular attention within ecological studies of freshwater zooplankton communities. In such communities, populations tend to fluctuate naturally in abundance throughout the year due to feedbacks between of population density and food availability (Sommer et al., 1986). Due to the importance of density and food abundance in regulating zooplankton populations, they present themselves as an ideal group of organisms in which to study indirect evolutionary responses to the introduction of novel predators, which may include shifts in the direction of trophic control and thus the strength of density dependence/resource limitation. Amongst zooplankton, cladocerans, and in particular *Daphnia*, have long served as model organisms in evolutionary biology. *Daphnia* present valuable assets such as clonal reproduction, short generation time, and are simple to maintain in laboratory conditions (Miner et al., 2012). Crucially, their ability to produce dormant resting eggs (ephippia) which can remain viable for centuries allows the application of “resurrection ecology” experiments, whereby phenotypes that have evolved under different environmental conditions can be compared in a common environment (Frisch et al., 2014).

Here, we leverage the invasion of the North European spiny water flea (*Bythotrephes longimanus*), a predator of herbivorous zooplankton, into Lake Kegonsa, Wisconsin, US (Barata et al., 2001; Walsh et al., 2016). Since *Bythotrephes* was first detected at high abundance in this lake in 2009, the biomass of one of its prey species, the cladoceran *Daphnia pulicaria*, has been reduced by up to 60% (Walsh et al., 2016). In a previous study of *D. pulicaria* from this lake, Landy et al. (2020) compared resurrected clones (i.e. genotypes) originating from prior to the invasion of *Bythotrephes* with those of contemporary (i.e., post-invasion) clones sourced directly from the lake, and provided evidence that the invasion has led to evolutionary change in a suite of life history and behavioral traits. Specifically, they demonstrated that invasion by *Bythotrephes* was associated with evolved reductions in size at maturity and fecundity. The evolutionary changes in these traits are hypothesized to have caused a decrease in the intrinsic rate of population growth of this species post-invasion. However, the measurements of body size and fecundity were obtained in ecologically benign conditions (i.e. under *ad libitum* food and no density dependence). In the current study, we follow up these findings and use population growth experiments, where density dependence and associated ecological effects such as resource limitation can manifest, to determine if evolutionary change in prey phenotypes can translate into evolved differences in population dynamics.

## Materials and Methods

### Study animals and husbandry

Sediment cores containing ephippia of *Daphnia pulicaria* and live individuals were collected from Lake Kegonsa, Wisconsin, US (42.96°, −89.31°) as described by Landy et al. (2020) in February 2018 and June 2019, respectively. Cores were ^210^Pb dated at the National Lacustrine Core Facility at the University of Minnesota, and ephippia from pre- and *post-Bythotrephes* invasion were transported to the University of Texas at Arlington for hatching. For this study, eight pre-invasion and eleven post-invasion clones were used (Table 1, Supplementary material). Hatched individuals from ephippia (representing all pre-invasion and three post-invasion clones, hatched during March 2019) and live-collected individuals (representing post-invasion individuals) were first kept at a 14L:10D photoperiod at 16°C in 90ml COMBO medium (Kilham et al., 1998) and fed non-limiting supply of green algae (*Scenedesmus obliquus*, ~1.0mg C L^-1^ day^-1^). In December 2019 live individuals of each clone were transported to the Norwegian University of Science and Technology and were subsequently kept at 17°C (photoperiod 16L:8D) in ADaM medium (Klüttgen et al., 1994, SeO2 concentration reduced by 50%, sea-salt increased to 1.23 g/L) and fed non-limiting supply of Shellfish Diet 1800 (Reed Mariculture Inc, Campbell, CA, USA) until the onset of the experiment. The same medium, food, temperature and light regime was used throughout the rest of the study.

**Table 1:** AICc comparisons of candidate models explaining variation in population abundance of experimental populations of *Daphnia pulicaria* originating from Lake Kegonsa. Separate models are fitted to numerical population abundance and biomass population abundance as dependent variables, and invasion history (whether the population originates from a period before or after invasion by the predatory zooplankton species *Bythotrephes longimanus*) as a fixed factor. Clone ID, population number and census number are included as random intercepts in all models.

|  | **K** | **AIC<sub>c</sub>** | **ΔAIC<sub>c</sub>** | **w<sub>i</sub>** |
| --- | --- | --- | --- | --- |
| **Numerical population abundance** |  |  |  |  |
| Invasion history | 6 | 14818.48 | 0.00 | 0.89 |
| ~1 | 5 | 14822.71 | 4.22 | 0.11 |
| **Biomass population abundance** |  |  |  |  |
| Invasion history | 6 | 4371.21 | 0.00 | 0.88 |
| ~1 | 5 | 4375.11 | 3.90 | 0.12 |

For each clone (n=19), 5-10 adult individuals were randomly chosen from stock cultures and placed in separate 1L glass beakers (1 clone per beaker) where they were fed 3.0ml shellfish diet every three days. When several egg-bearing individuals were identified in each beaker, these were isolated by removing all others. Beakers with egg-bearing individuals were checked for neonates every 24h and each neonate found within this period was individually transferred to a plastic container (C577-120W) containing 100ml of ADaM. Newborn individuals that died within 6 days were replaced using the same method. In total, 10 newborns were selected from each clone over a span of seven days, yielding 190 populations that originally consisted of a single individual (10 individuals per clone x 19 clones, 8 pre-invasion, 11 post-invasion). When an individual died after the 7^th^ day, it was reported as dead and not replaced (n = 11). In total, 21 pre-invasion populations (23.3%) and 9 post-invasion (9.0%) populations went extinct during the experiment, and 66.7% of these extinctions occurred within the first 10 days of the experiment. The plastic containers were stored in 2 Memmert Peltier cooled incubator IPP 260plus (Memmert, Germany) climate cabinets at 20.0 °C (photoperiod 16L:8D). 1.0 mL of shellfish diet was added every second day and medium was changed every eight days. Container placement in the climate cabinets was changed haphazardly every two days, after feeding. All populations time series were run in parallel during March-May 2020.

### Measuring population growth

To obtain data on population growth, video recordings were made of each population every 8 days (with one exception due to covid-19 regulations) for a period of three months, starting 11 days after the start of the population growth experiment, creating a total of 10 censuses. To prepare each population for filming, the content of each container was poured into a second plastic container fitted with a fine mesh sieve to remove most of the medium. *Daphnia* were subsequently flushed (with ADaM) from the sieve with the help of a spraying bottle into a transparent square tray (9.0×9.0cm). Large pieces of algae were removed from the tray by pipetting. The empty plastic container was also sprayed with ADaM to remove attached algae and leftover *Daphnia*. This content was poured into a second tray and if *Daphnia* were found, these were moved with a pipette to the main tray.

The transparent tray was put on a LED light board (Huion A4 LED light pad, set to maximum intensity) in a dark room directly under the camera (Basler aCA1300-60gm, fitted with 5–50-mm, F1.4, CS mount lens) and filmed for 10-14 seconds. We waited for any movement of the medium to stop before starting each recording. The contents of the tray were then emptied back in the original plastic container. This procedure was repeated for each population. After finishing, each container was filled to 100mL with ADaM and 1.5 mL shellfish diet was added. The larger ration during this event than during regular feeding was done to compensate for the complete removal of all algae and bacteria during medium change. The procedure described above produced a total of 1657 videos that were subsequently analyzed to obtain data on population sizes. The first 39 frames from each video were extracted as images using a custom Matlab script. The images were then analyzed using the R package *trackdem* (Bruijning et al., 2018). *trackdem* estimates the number of live individuals (based on movement and contrast) and their size (based on number of pixels) in a sequence of images. Dry mass (mg) of each individual was calculated as −0.00635 + 0.00100 x pixel number, based on a regression developed for *D. magna* (Fossen et al., 2021). Population sizes from 18 sample videos selected from throughout the experimental period were counted manually and optimal *trackdem* settings were found by regressing these against estimated counts from *trackdem*. For the optimal settings (Supplementary material), the fitted regression was *trackdem* count = 0.08 + 0.96 x manual count (R^2^ = 0.93, n = 18).

Daily population growth rates, *G*, were calculated as *G* = log_e_(*N*_t+1_/ *N*_t_)/*d*, where *N*_t_ and *N*_t+1_ is the population abundance (measured either in number of individuals or total biomass) at two consecutive censuses, and *d* is the duration of time in days between the censuses. Growth rate was not calculated between two censuses if the population went extinct. Outliers were identified by inspecting residuals from a linear regression between population count and growth rate. Manual inspection of videos with high residual values identified 16 instances where *trackdem* estimates of population size were unexpectedly large. These values were corrected by conducting manual counts from the respective videos. All of these cases were caused by a high abundance of moving algae, mistakenly counted as *Daphnia* by *trackdem*. Manual inspection of videos producing negative outliers were confirmed to be valid and caused by rapid population declines.

### Choice of population dynamics model and calculation of *r*

Inspection of population growth rate data (both for numerical and biomass growth) revealed strong non-linearity in the density dependence. Thus, the population dynamics is best described by the theta-logistic model *G* = *r*(1-(*N*_t_/*K*)^*θ*^) where *r* is the intrinsic population growth rate, *K* is the carrying capacity, and *θ* determines the shape of the density dependence. One concern when fitting this model to data is that different combinations of *r* and *θ* can produce model fits of similar likelihood, potentially resulting in ecologically unrealistic estimates of both parameters (Clark et al. 2010). We therefore took advantage of the experimental design, whereby each population was started with a single neonate, which allowed us to obtain direct observations of population growth under low density. We based this calculation of *r* for each population on the observed population growth rate from the start of the experiment until the second census (i.e. *r* = loge(*N*_2_/ *N*_0_)/*d*). This duration (i.e. 18 days) is similar to the typically used time span of 21 days used in estimation of *r* from life table data in *Daphnia*. This was done both for numerical and biomass growth.

### Statistical analyses

All statistical analyses were conducted in R v.4.1.0 (R Core Team, 2021). To test for an overall difference in numerical abundance and population biomass throughout the experiment, linear mixed-effect models were fitted using the function *lmer* in the package *lme4* (Bates et al., 2015). The full model contained the fixed effect of invasion history (pre- vs. post-invasion type). Random effects were clone ID, experimental population number nested within clone ID, and a crossed random effect of census number. The random effect of census number was chosen as we were not interested in explicitly modelling the temporal trends in population abundance, but had to account for the multiple observations per population.

We tested for an effect of invasion history on the observed values of *r* (calculated early in the sequence of population growth and thus under low density, see above) by fitting a linear mixed model to these data, again using *lmer*, including a random effect of clone ID, and comparing this against a simpler model containing only the random effect using the Akaike information criterion corrected for small sample sizes (*AICcmodavg*, Mazerolle, 2020). This was done both for numerical and biomass growth rate.

Next, to test for an effect of invasion history on *K* we fitted non-linear mixed effect models representing the theta-logistic model to the population growth time series. This was done for the data following the first two censuses, i.e. after the period that had been used to calculate *r*. Again, this was done separately for numerical and biomass growth rates. In the full model, population growth rate over a given period (between two consecutive censuses) was modelled as a function of the observed value of *r* for that population and its population size at the start of that period while estimating values of *K* that depended on invasion history, and a single value of *θ* common to all populations. The model included random effects of clone ID and population nested within clone ID on *K*. The model was fitted using the function *nlme* in the package *nlme* (Pinheiro et al. 2021). This full model was compared to a simpler one where *K* was common to all populations independent of invasion history, again using the Akaike information criterion corrected for small sample sizes. For all analyses assumptions of normality and homogeneity of residuals were satisfied. All figures were made using the package *ggplot2* (Wickham, 2016).

## Results

Population size trends for both pre- and post-invasion populations showed a period of high population growth up to a peak level, followed by a slower decline (Fig. 1). This was true both for numerical abundance and biomass. For both measures of population size, post-invasion clones maintained higher values than pre-invasion clones throughout the experimental period (Fig. 1, Table 1). Model parameter estimates suggest that post-invasion clones attained an overall increase relative to pre-invasive clones in numerical abundance by 28%, and in biomass abundance by 33% (Table 2).

**Figure 1:**
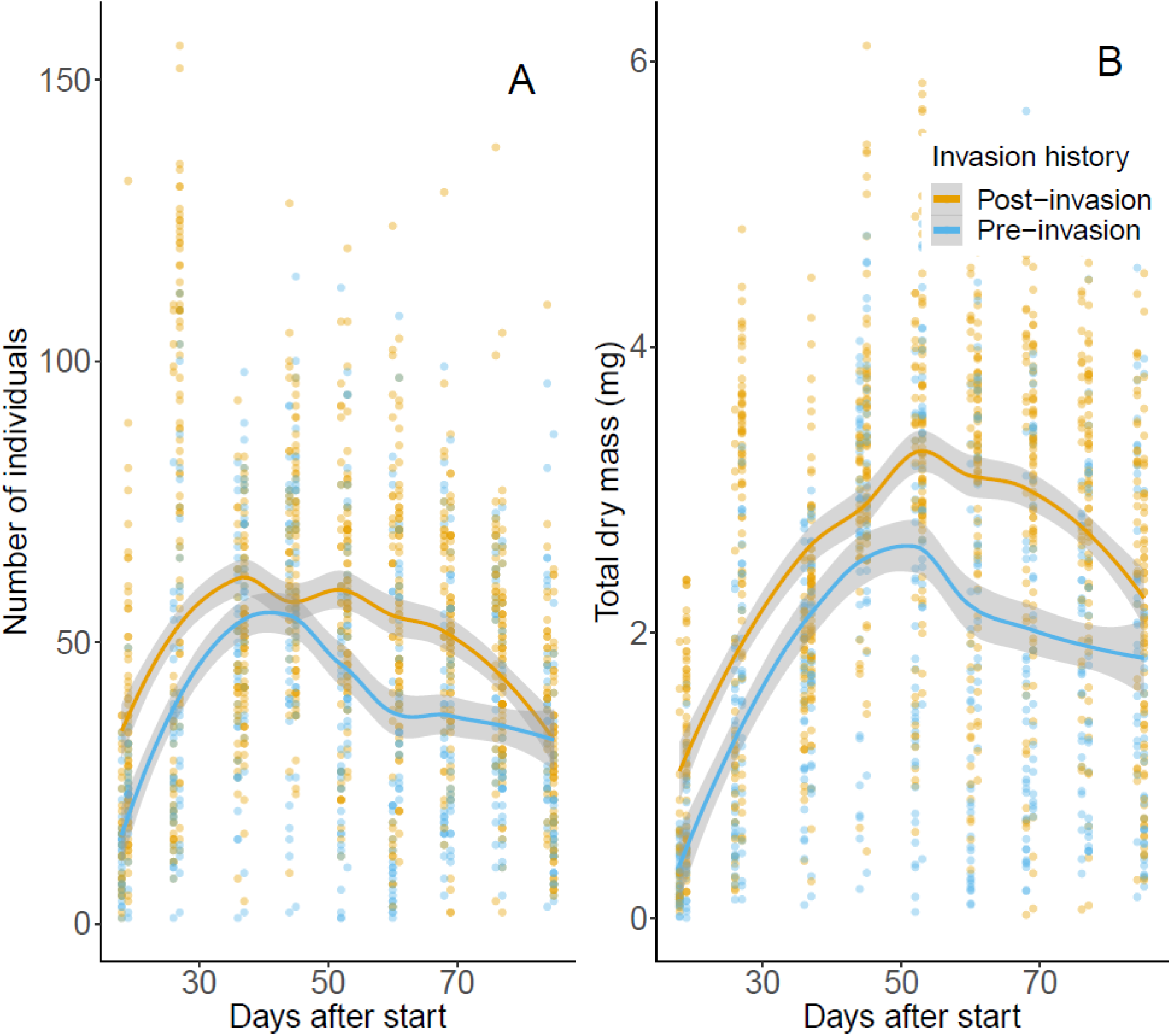
Population size trends in terms of (A) number of individuals and (B) total dry mass for experimental populations of *Daphnia pulicaria* originating from Lake Kegonsa. Pre- and post-invasion populations consist of clones sourced from before and after *Bythotrephes longimanus* invasion in 2009, respectively. Lines are fitted with geom_smooth (method=‘loess’) and shaded areas represent 95% CI.

**Table 2:** Summary of the best fitting linear mixed-effect models (fitted using REML) estimating the effect of invasion history on numerical and biomass population abundance of *Daphnia pulicaria* originating from Lake Kegonsa (Table 1). Model estimates for pre-invasion populations are presented relative to the respective estimates for post-invasion populations. Post- and pre-invasion populations consist of clones originating from after and before *Bythotrephes longimanus* invasion, respectively.

|  | Estimate | SE |
| --- | --- | --- |
| **Numerical population abundance** |  |  |
| **Fixed effects** |  |  |
| Intercept | 45.66 | 5.88 |
| Period (pre-invasion) | -10.00 | 3.77 |
| **Random effects (SD)** |  |  |
| Clone ID | 7.43 |  |
| Population:Clone ID | 6.04 |  |
| Census number | 16.94 |  |
| **Biomass population abundance** |  |  |
| **Fixed effects** |  |  |
| Intercept | 2.26 | 0.33 |
| Period (pre-invasion) | -0.56 | 0.22 |
| **Random effects (SD)** |  |  |
| Clone ID | 0.44 |  |
| Population:Clone ID | 0.38 |  |
| Census number | 0.94 |  |

Observed values of intrinsic population growth rate also tended to depend on invasion history (Table 3). The evidence for such an effect was strongest for biomass (Table 3), but models containing an effect of invasion history suggested higher intrinsic population growth rate in post-invasive clones than in pre-invasive clones for both numerical and biomass growth (Table 4). The estimated increase in intrinsic population growth rate based on numerical and biomass data were 23 and 15%, respectively (Table 4).

**Table 3:** AICc comparisons of candidate models explaining variation in observed intrinsic population growth rate (*r*) of experimental populations of *Daphnia pulicaria* originating from Lake Kegonsa. Separate linear mixed effects models are fitted to numerical population growth rate and biomass population growth rate as dependent variables. Full models include effects of invasion history (whether the population originates from a period before or after invasion by the predatory zooplankton species *Bythotrephes longimanus*), with Clone ID as a random effect.

|  | **k** | **AICc** | **ΔAICc** | **w<sub>i</sub>** |
| --- | --- | --- | --- | --- |
| **Numerical population growth rate** |  |  |  |  |
| $r \sim$ invasion history | 4 | -7147.33 | 0.00 | 0.64 |
| $r \sim 1$ | 3 | -7146.18 | 1.15 | 0.36 |
| **Biomass population growth rate** |  |  |  |  |
| $r \sim$ invasion history | 4 | -6626.54 | 0.00 | 0.75 |
| $r \sim 1$ | 3 | -6624.34 | 2.19 | 0.25 |

**Table 4:** Parameter estimates (obtained using REML) of the best fitting models describing the variation in observed intrinsic population growth rate (*r*) of *Daphnia pulicaria* originating from Lake Kegonsa (Table 3). Post-and pre-invasion populations consist of clones originating from after and before *Bythotrephes longimanus* invasion, respectively.

|  | Estimate | SE |
| --- | --- | --- |
| **Numerical population growth rate** |  |  |
| **Fixed effects** |  |  |
| $r$ post-invasion (intercept) | 0.16 | 0.01 |
| $r$ pre-invasion (difference) | -0.03 | 0.02 |
| **Random effects (SD)** |  |  |
| Clone ID | 0.04 |  |
| **Biomass population growth rate** |  |  |
| **Fixed effects** |  |  |
| $r$ post-invasion (intercept) | 0.31 | 0.01 |
| $r$ pre-invasion (difference) | -0.04 | 0.02 |
| **Random effects (SD)** |  |  |
| Clone ID | 0.04 |  |

The difference in population dynamics between pre- and post-invasion clones was also reflected in the comparisons of theta-logistic models of population growth rates. For both numerical and biomass data, models containing an effect of invasion history on the carrying capacity received more support than models without this term (Table 5). The strength of evidence for such an effect was particularly pronounced for numerical data. For both these analyses, the best models predicted a higher carrying capacity for post-invasion clones compared to for the pre-invasion clones (Table 6, Fig. 2). The estimated increase in carrying capacity based on numerical and biomass data were 27 and 25%, respectively (Table 6).

**Figure 2:**
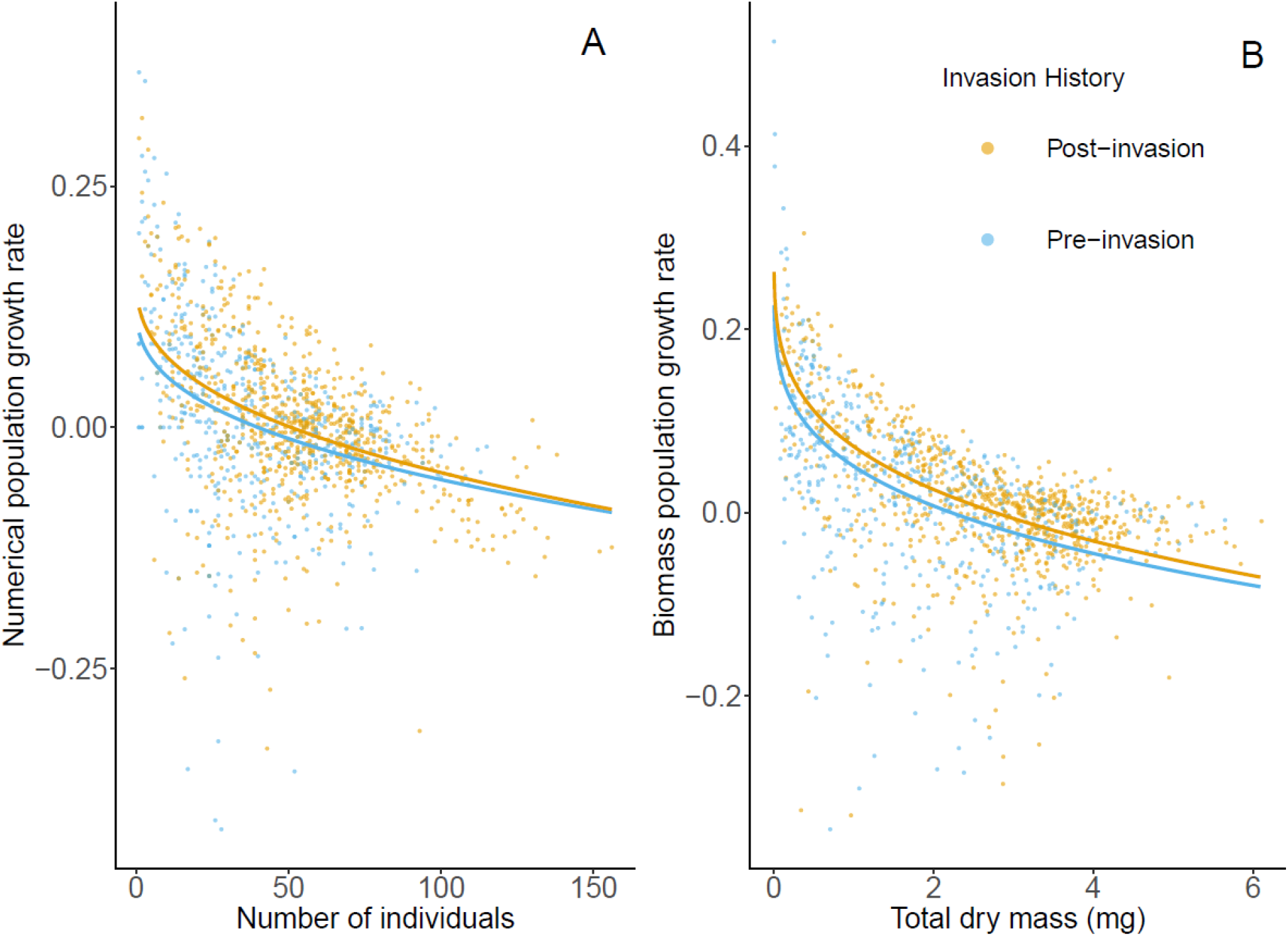
Growth rate in terms of (A) number of individuals and (B) total dry mass for pre- and post-invasion populations of *Daphnia pulicaria* originating from Lake Kegonsa. Pre- and post-invasion populations consist of clones sourced from before and after *Bythotrephes longimanus* invasion in 2009, respectively. Regression lines give predictions from theta-logistic models with parameter estimates from Table 4 (for *r*) and Table 6 (for *K* and *θ*).

**Table 5:** AICc comparisons of candidate models explaining variation in population growth rate of experimental populations of *Daphniapulicaria* originating from Lake Kegonsa. Separate theta-logistic models are fitted to numerical population growth rate and biomass population growth rate as dependent variables. Full models include effects of invasion history (whether the population originates from a period before or after invasion by the predatory zooplankton species *Bythotrephes longimanus*) on *K*. Clone ID and population (nested within Clone ID) are included as random effects on *K* in all models.

|  | **k** | **AICc** | **ΔAICc** | **w<sub>i</sub>** |
| --- | --- | --- | --- | --- |
| **Numerical population growth rate** |  |  |  |  |
| $K \sim$ invasion history | 6 | -3010.67 | 0.00 | 0.95 |
| $K \sim 1$ | 5 | -3004.92 | 5.75 | 0.05 |
| **Biomass population growth rate** |  |  |  |  |
| $K \sim$ invasion history | 6 | -3314.14 | 0.00 | 0.67 |
| $K \sim 1$ | 5 | -3312.70 | 1.43 | 0.33 |

**Table 6:** Parameter estimates (obtained using REML) for *K*, *θ* and random effects of *K* of the best fitting theta-logistic models describing the population dynamics of *Daphnia pulicaria* originating from Lake Kegonsa (Table 1). Post-and pre-invasion populations consist of clones originating from after and before *Bythotrephes longimanus* invasion, respectively. t- and p-values refer to the difference in estimated *K* for the two invasion histories.

|  | Estimate | SE |
| --- | --- | --- |
| **Numerical population growth rate** |  |  |
| **Fixed effects** |  |  |
| $K$ post-invasion (intercept) | 50.88 | 2.11 |
| $K$ pre-invasion (difference) | -10.89 | 3.08 |
| $\theta$ | 0.38 | 0.03 |
| **Random effects (SD)** |  |  |
| Clone ID | 0.0033 |  |
| Population:Clone ID | 0.0011 |  |
| **Biomass population growth rate** |  |  |
| **Fixed effects** |  |  |
| $K$ post-invasion (intercept) | 2.77 | 0.18 |
| $K$ pre-invasion (difference) | -0.55 | 0.27 |
| $\theta$ | 0.26 | 0.01 |
| **Random effects (SD)** |  |  |
| Clone ID | 0.5069 |  |
| Population:Clone ID | 0.0001 |  |

## Discussion

We leveraged a well-documented invasive event by a predatory species (*Bythotrephes longimanus*) in combination with a resurrection ecology approach to investigate evolutionary changes in population dynamics of their main prey species (*Daphnia pulicaria*). This allowed us test for an association between a change in predation levels and evolution of prey population dynamics in a natural population. Post-invasion genotypes were able to maintain a higher population abundance throughout our experiment, both in terms of the number of individuals and population biomass. Estimation of population dynamics parameters suggested that this was achieved through an increase in intrinsic population growth rate *r* (strong evidence for biomass data) as well as carrying capacity *K* (strong evidence for numerical data). For the current study we had prior information on evolutionary change in individual traits in response to the predator invasion. Specifically, for traits that can be directly used to predict effects on population dynamics, Landy et al. (2020) found that post-invasion clones had a reduced size at maturity and reduced fecundity compared to pre-invasion clones, but found no significant difference in age at maturity. Based on this one might predict a reduced intrinsic rate of increase in post-invasive clones. Yet, we observed the opposite in our study. This indicates that care should be taken when extrapolating life history traits measured in isolated individuals under benign laboratory conditions to estimates of population dynamics. One reason for this is that unmeasured traits that may be genetically correlated with those that are measured may also influence population dynamics. For example, if offspring quality and survival is negatively correlated with fecundity (Mappes & Kostela, 2004), these two traits may counteract each other in terms of effects on population growth. Similarly, as highlighted by the pace-of-life syndrome literature (Wright et al., 2019), life history traits may be correlated with behavioral traits that in turn influence social interactions and hence the intrinsic rate of increase as well as responses to increasing population density. The results from the current study support the conclusion from these considerations; studies using actual population growth experiments may be required to understand how ecological interactions shape evolutionary change in population dynamics.

Although we are not aware of previous studies that have quantified the evolutionary effect of predation level on the evolved dynamics of wild prey populations, a series of chemostat experiments have addressed the role of evolution in shaping the population dynamical response of planktonic algae (*Chlorella vulgaris*) to rotifer (*Brachionus calyciflorus*) predation. These studies have demonstrated that the patterns of predator-prey cycles are influenced by prey evolution (Shertzer et al., 2002, Yoshida et al., 2003). This system has also been used to test for an evolutionary trade-off between algal population growth and predator defense (Yoshida et al., 2004). Algae were allowed to evolve in the presence and absence of the rotifer, after which their growth rates were measured at different nutrient-levels in the absence of predation. The results showed that algae that had evolved under predation exposure had a lower population growth rate than those having evolved in absence of predation, but only at the most limiting nutrient-level. When nutrients were more abundant no such difference was observed. Thus, the authors conclude that the algae face a trade-off between competitive ability and predator resistance, and additional lines of evidence suggest that this is due to selective grazing by rotifers on larger cells, rather than indirect selective effects (Yoshida et al., 2004).

In the current study we found no evidence for an interaction between predation history and food availability in determining the rate of population growth as observed in the rotifer/algae experiments. Rather, clones that had evolved in the presence of the predator were able to maintain an overall larger population abundance throughout the experiment (and thus in response to declining levels of food as population density increased) in the absence of the predator. Furthermore, models that contained an effect of invasion history consistently suggested elevated values of both *r* and *K* in post-invasion clones compared to pre-invasion clones, independent of measurement type (numerical or biomass population dynamics). Thus, our results also fail to provide support for an evolutionary trade-off between the intrinsic rate of increase and strength of density dependence that has been predicted on theoretical grounds (i.e. *r* vs. *K*, Boyce, 1984). This may indicate that evolutionary responses to invasive predators in natural systems can be more complex than what can be extrapolated from experimental evolution studies or predicted from theory. We propose two explanations that may have played a role in creating such complexities for the predator-prey system investigated here. First, as for most zooplankton, both species show extensive seasonal dynamics in abundance, but they are not synchronous. Data from nearby Lake Mendota, another lake having experienced a recent *Bythotrephes* invasion, show that whereas *D. pulicaria* population abundance peaks during early summer (May/June), *Bythotrephes* populations have a slower growth rate and continue to grow from April until they peak in October (Walsh et al., 2016). Thus, the invasion and associated heavy predation towards the last part of the growth season may have strengthened selection for rapid growth, favoring clones that reach high abundance prior to the onset of high predation rates. We do not have data on the timing of resting egg production in Lake Kegonsa, which would indicate the importance of reaching high frequency early in the season. However, sampling in late September 2018 showed that *D. pulicaria* was largely absent by that time (M. Walsh, unpublished data), suggesting that the largest contribution to the resting egg bank may occur during summer. Another potentially complicating factor is the propensity for *D. pulicaria* to migrate vertically in the presence of a predator, whereby they move to deeper parts of the lake during the day to avoid predation. Indeed, Landy et al. (2020) found evolution towards reduced positive phototaxis in post-invasive clones from Lake Kegonsa, suggesting an increased propensity to undertake such migrations. *Daphnia* feed on phytoplankton, which tend to congregate at the lake surface. Thus, life in deeper water also means living in a more resource limited environment (Cousyn et al., 2001; Pangle & Peacor, 2006; Pijanowska et al., 1993). Previous studies have shown that organisms living at different resource levels may evolve adaptations to this, such that when reared in a common environment, those from more food-restricted environments actually grow faster (Arendt & Wilson 1999; Iraeta et al. 2013). Thus, our observation that post-invasive clones outperform pre-invasive clones may be an example of such countergradient variation (Conover & Schultz 1995), where they have evolved physiological adaptations that increase population growth under a given level of food abundance. In a recent study of *D. pulicaria* from Lake Mendota, Rani et al. (2021) found that post-invasive clones had a reduced metabolic rate compared to pre-invasion clones, which may represent one such physiological adaptation to a cooler resource deficient environment. Furthermore, Einum et al. (2019) found that variation in somatic growth rate among clones of *D. magna* was best explained by clone-specific food consumption expressed relative to their rate of energy use. Thus, if post-invasion clones have a reduced rate of metabolism but do not moderate food consumption when reared in a common environment (as in the current study), this could be expected to result in increased somatic growth rate and may thus explain the higher population growth rate.

To conclude, the current study demonstrates an evolutionary shift in the population dynamics of *D. pulicaria* in parallel with an increase in predation brought about by invasion of the predatory zooplankton *Bythotrephes*. To our knowledge, this is the first empirical study that directly demonstrates this by obtaining time series of population growth in a common environment and comparing genotypes that come from the same population but have contrasting histories of exposure to predation. The observed shift in population dynamics may be related to selection for reduced predator exposure, either temporally or spatially, and suggest that complexities in ecological interactions may make it challenging to apply knowledge from experimental evolution studies and theory to predict the evolutionary responses of population dynamics to changes in predation pressure in natural systems.

## Data Archiving Statement

Data for this study are available at: *to be completed after manuscript is accepted for publication*.

## Acknowledgements

The study was funded through the Research Council of Norway’s (RCN) project numbers 230482, 244046 and 223257. We thank V Rani for lab assistance.

